# CBM KG: A Comorbidity-Centric Knowledge Graph Uncovering Causal Pathomechanisms Between COVID-19 and Neurodegenerative Diseases

**DOI:** 10.1101/2025.08.18.670790

**Authors:** Heval Atas Guvenilir, Priya Sethumadhavan, Abish Kaladharan, Renata Vieira de Sa, Adithya Sridhar, Undine Haferkamp, Ole Pless, Andrea Martí-Sarrias, Sandra Acosta, Tamas Letoha, Anett Hudak, Jochen Ohnmacht, Feng Q. Hefeng, Martin Hofmann-Apitius, Alpha Tom Kodamullil

## Abstract

**Summary:** COVID-19 is increasingly recognized as a potential trigger or accelerator of neurodegenerative diseases such as Alzheimer’s and Parkinson’s. To systematically explore the putative molecular and clinical associations between them, we present CBM KG (Causal Biological Mechanisms Knowledge Graph)—a manually curated, comorbidity-centric resource developed within the EU-funded COMMUTE project. CBM KG integrates over 2,800 cause-and-effect or correlative relationships from 63 peer-reviewed publications, highlighting key mechanisms such as viral entry routes, blood-brain barrier alteration, microglial activation, neuroinflammation, and APOE ε4-associated susceptibility. Each relationship in the graph is fully traceable to its source evidence, ensuring transparency and reproducibility. Unlike general-purpose or single disease-focused knowledge graphs, CBM KG is specifically designed to represent causal biological mechanisms spanning both infectious and neurodegenerative processes. By encoding directional, cause-and-effect relationships, it supports the interpretation of clinical co-occurrences through plausible mechanistic links between overlapping disease pathways, offering high-resolution insights at both molecular and clinical levels.

**Availability and implementation:** The BEL files, Neo4j database, and Cytoscape visualization files are publicly available at: https://github.com/SCAI-BIO/CBM-Comorbidity-KG.

## Introduction

The COVID-19 pandemic, caused by SARS-CoV-2, has presented a global health crisis with consequences extending far beyond the acute infection phase. Emerging evidence highlights its role in triggering or exacerbating chronic comorbidities, including neurodegenerative diseases (NDDs) such as Alzheimer’s disease (AD) and Parkinson’s disease (PD). Epidemiological studies have reported an increased incidence of AD and PD diagnoses, as well as accelerated symptom progression in post-COVID-19 patients, suggesting overlapping pathophysiology mechanisms (Taquet et al., 2021; Li et al., 2022). Understanding the complex interplay between SARS-CoV-2 infection and these chronic conditions is therefore crucial to estimate the long-term consequences of the pandemic.

Comorbidity refers to the co-occurrence of distinct clinical conditions within the same individual (Valderas et al., 2009), often identified through observational studies without mechanistic explanation. Knowledge graphs (KGs) provide a powerful framework to represent such complex disease knowledge in a computable format, facilitating the visualization and traversal of interconnected relationships (Himmelstein et al., 2017; Chandak et al., 2023). While general-purpose KGs offer broad coverage of biological entities and disease-specific KGs provide detailed insights into single conditions, neither approach alone is optimized to capture the complex interplay between distinct disease processes.

Within the broader context of the EU-funded COMMUTE project (www.commute-project.eu), which integrates data-driven and knowledge-based approaches to investigate the long-term neurological sequelae of COVID-19, we developed the Causal Biological Mechanisms Knowledge Graph (CBM KG) as a manually curated, comorbidity-centric, cause-and-effect KG linking SARS-CoV-2 infection to AD and PD. Here, comorbidity-centric denotes the targeted curation of literature addressing both conditions in a shared context. Encoded in Biological Expression Language (BEL; https://bel.bio/) and implemented in a Neo4j database, the CBM KG comprises over 2,800 causal or correlative relationships extracted from 63 peer-reviewed publications. It captures a diverse spectrum of multi-level interactions spanning viral entry mechanisms, neuroinflammatory cascades, blood-brain barrier alteration, and genetic risk factors, as well as convergent pathways such as oxidative stress and protein aggregation. By integrating not only proteins and genes, but also protein/gene variants, fragments, complexes, and post-translational modifications, CBM KG provides high resolution molecular level information, enabling the construction of more informative causal chains connecting molecular mechanisms into clinical outcomes. It also stores the relevant evidence statements and associated PMIDs for each relationship as edge properties, enabling traceability to the original publication and enhancing the reliability of the data. To our knowledge, CBM KG is the first manually curated, comorbidity-centric KG specifically designed to model molecular level cause-and-effect relationships between COVID-19 and neurodegeneration, addressing a critical need in post-pandemic healthcare. This level of refinement is essential for generating actionable hypotheses, prioritizing biomarker candidates, and guiding therapeutic strategies in complex, multifactorial disease contexts, capabilities that extend beyond those of conventional knowledge graphs. CBM KG is publicly accessible at https://github.com/SCAI-BIO/CBM-Comorbidity-KG for querying, extension, and functional interpretation of experiments aimed at unraveling the molecular and cellular basis of the putative comorbidity between COVID-19 and neurodegenerative diseases.

## Approach

### Data Curation and BEL Encoding

Based on expert opinion contributed by COMMUTE biomedical and clinical research experts, we created a highly relevant literature corpus focusing on the complex relationship between COVID-19 (SARS-CoV-2) and neurodegeneration (particularly Alzheimer’s disease and Parkinsonism). Then, curation experts at Causality BioModels (CBM), a curation spin-out of Fraunhofer SCAI (https://www.causalitybiomodels.com), analysed these 63 full-text publications (50 research articles, 13 reviews) and extracted approximately 2,000 unique BEL statements yielding over 2,800 relationships (subject-predicate-object triples). These triples encode cause-and-effect (e.g., “increases”, “decreases”, “directly_increases”, “directly_decreases”) or correlative relationships (e.g., “positive_correlation”, “negative_correlation”) among various biological entities including genes, proteins, drugs, compounds, phenotypes, molecular and biological processes, pathologies and clinical endpoints, as well as variants, fragments, and complexes. To ensure semantic consistency and reusability, the extracted relationships were formalized using BEL (https://bel.bio/). BEL captures relationships in a structured manner, enabling computational reasoning and integration with existing BEL-based resources. We stored BEL triples and statements of each publication as a separate .bel file so that we maintain full provenance for each BEL triple.

### KG Construction, Visualization, and Analysis

Each BEL file was converted into JSON format and imported into a Neo4j (v5.20) graph database using the eBEL Python package (available at https://github.com/e-bel/ebel/tree/neo4j), ensuring harmonization of shared entities across separate JSON files. The resulting KG involves 2,424 nodes and 5,043 directed edges, enriched with diverse relationship types from the eBEL conversion process. For interactive exploration and visualization, the graph was then imported into Cytoscape (v3.10.2) (Shannon et al., 2003) via the Neo4j plugin, which retrieves network data through Cypher queries. Disease-related sub-networks were subsequently extracted using Cytoscape’s built-in functionalities, including filtering and selection tools.

## Results

The workflow proposed in this study transforms unstructured literature into a computable, knowledge-driven network for mechanistic exploration. Our analyses reveal potential mechanisms and dysregulated pathways linking COVID-19 and NDD progression. The knowledge graph (*See Figure 1b and Supplementary Table 1 “ST1” for the corresponding BEL triples and their supporting evidence statements related to the following mechanisms*) illustrates canonical viral entry mechanisms involving ACE2 receptors and TMPRSS2 protease (ST1-E1), which, while expressed at low levels across most brain regions, are reported to be upregulated in neurons and choroid plexus (ChP) epithelial cells following infection (ST1-E2 & E3). These observations, primarily derived from human pluripotent stem cell-derived brain organoid models, suggest that the ChP could represent a vulnerable interface through which SARS-CoV-2 may gain access to the central nervous system (CNS). Additionally, the graph captures alternative, ACE2-independent entry routes via receptors including DPP4 and BSG, particularly in astrocytes (ST1-E4), and NRP1, expressed across neurons and glial cells (ST1-E5 & E6), together suggesting broader viral tropism within the CNS. Once inside the CNS, the virus appears to compromise the integrity of the blood-brain barrier (BBB) (ST1-E7), facilitating the infiltration of inflammatory cytokines, immune cells, and viral particles into the brain (ST1-E8). The knowledge graph indicates that the immune response to SARS-CoV-2 is associated with elevated cytokine production (ST1-E9), which may drive microglial activation (ST1-E10) and chronic inflammation (ST1-E11), common processes in NDD pathophysiology. Activated microglia, in turn, releases pro-inflammatory mediators such as IL-6 and TNF (ST1-E11), further contributing to neuronal damage (ST1-E12). Notably, this neuroinflammatory cascade may occur even without direct viral invasion of the CNS (Matschke et al., 2020; Thakur et al., 2021), as systemic cytokine elevations in severe COVID-19 can impair the blood–brain barrier and indirectly trigger glial activation and neuroinflammation (Vanderheiden and Klein, 2022). CBM KG also suggests that SARS-CoV-2 infection may promote the accumulation of alpha-synuclein through inflammatory responses (ST1-E13), contributing to Lewy body formation (ST1-E14), and may trigger an immune response that releases reactive oxygen species (ST1-E10) and nitric oxide (ST1-E15). Collectively, these events impair neuronal function and survival, ultimately leading to progressive dopaminergic neuron loss observed in PD (ST1-E16 & E17). Additionally, CBM KG captures genetic factors, such as the AD risk allele APOE e4, linked to increased susceptibility to severe COVID-19 outcomes (ST1-E18). These findings, supported by the broader literature incorporated within the CBM KG, underscore the complex interplay and potential comorbidity between SARS-CoV-2 infection and neurodegenerative processes, highlighting the need for further investigation.

**Figure 1.**
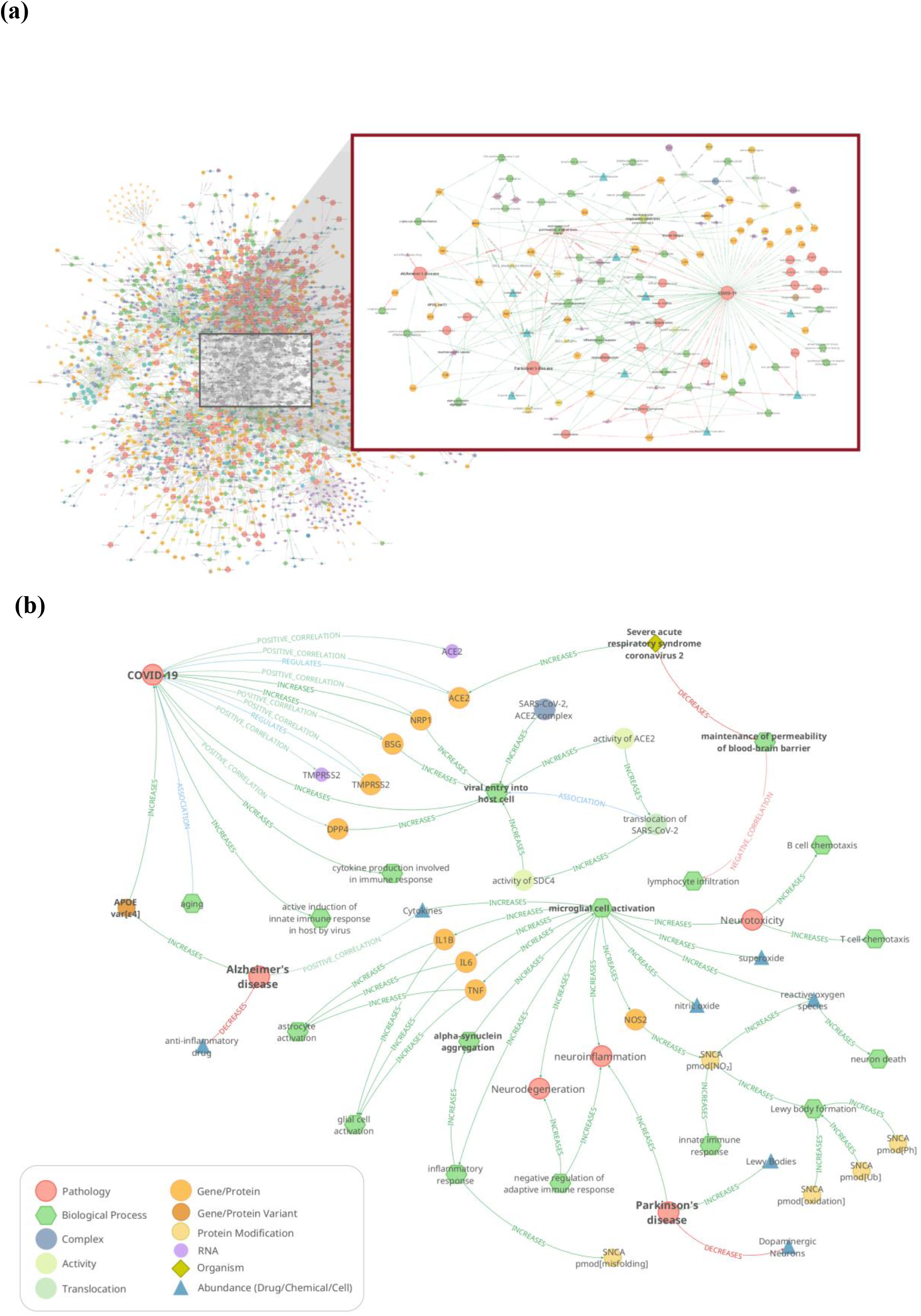
**(a)** Overview of the CBM KG highlighting putative comorbidity mechanisms between COVID-19 and neurodegenerative diseases. **(b)** Subgraph illustrating plausible causal cascades connecting SARS-CoV-2 infection to downstream molecular mechanisms implicated in the progression of AD and PD.

## Discussion

CBM KG addresses a critical need by providing mechanistic insights into the interplay between SARS-CoV-2 infection and neurodegenerative diseases. Unlike traditional KGs, it encodes directional, cause-and-effect relationships that span both conditions, offering enhanced interpretability for comorbid conditions. This refined framework facilitates the exploration of shared pathophysiology mechanisms and supports the identification of potential therapeutic interventions at the intersection of viral infection and neurodegeneration. Leveraging BEL to encode cause-and-effect relationships and structuring them within a Neo4j-backed KG, we provide a core resource for hypothesis generation, functional interpretation of experimental data and therapeutic target discovery. Beyond genes, proteins, and clinical phenotypes, CBM KG integrates additional layers of biological information - such as genetic variant data, protein fragments, and complexes - that enrich the network’s contextual relevance. In contrast to general-purpose or single-disease-specific graphs, our comorbidity-centric approach delivers a way more specific representation of overlapping disease mechanisms, which is essential for tackling the multifaceted clinical challenges posed by COVID-19.

Future work can extend this study by integrating experimental data - such as gene expression profiles, single nucleotide polymorphism (SNP) data, and other omics datasets - to validate and refine the causal pathways identified by CBM KG. This integrative validation will not only enhance the robustness of the KG but also increase its translational potential for precision medicine. Additionally, by incorporating advanced AI methodologies, the framework could be further expanded to identify novel therapeutic targets, profile high-risk patient cohorts, and devise effective management strategies for neurodegenerative conditions.

