## Supplementary Table 1 for "CBM KG: A Comorbidity-Centric Knowledge Graph Uncovering Causal Pathomechanisms Between COVID-19 and Neurodegenerative Diseases"

**Supplementary Table 1.** Source PMIDs and supporting evidence for corresponding BEL triples.

| Entry ID (BEL line reference) | PMID (Source BEL file) | BEL Triple(s) | Evidence Statement |
|---|---|---|---|
| E1 (267-269) | 37798476 | p(HGNC:ACE2) regulates path(DO:"COVID-19")<br><br>p(HGNC:TMPRSS2) regulates path(DO:"COVID-19") | "ACE2 and cellular proteases, such as TMPRSS2, have been identified as receptors and priming factors for SARS-CoV-2." |
| E2 (161-169) | 33631122 | path(DO:"COVID-19") pos p(HGNC:ACE2)<br>path(DO:"COVID-19") pos p(HGNC:TMPRSS2) | "Our results showed that infection was more efficient in neurons than in NPCs (Figures 3F and 3G), an observation that correlates with the levels of ACE2 and TMPRSS2 expression in neurons and NPCs (Figures 3B and 3C)." |
| E3 (175-176) | 33113348 | a(TAX:"Severe acute respiratory syndrome coronavirus 2") increases p(HGNC:ACE2) | “Again, this infection matched ACE2 expression, with infected cells co-staining for the receptor (Figure 4A) in the ChP epithelium, whereas stromal cells of the ChP were uninfected (Figure S4A).” |
| E4 (121-125) | 37458167 | SET MeSHAnatomy = "Astrocytes"<br>p(HGNC:BSG) -> bp(GO:"viral entry into host cell")<br>p(HGNC:DPP4) -> bp(GO:"viral entry into host cell") | "Recently, the extracellular proteases BSG and DPP4 have also been identified as additional entry receptors for SARS-CoV-2 on astrocytes, providing new detail on the neuropathogenesis of COVID-19 (Andrews et al., 2022; Wang et al., 2020) (Fig. 2)." |
| E5 (116-119) | 37458167 | SET MeSHAnatomy = {"Brain","Neurons", "Astrocytes", "Microglia"}<br>p(HGNC:NRP1) -> path(DO:"COVID-19") | "Studies based on publicly available transcriptome databases revealed that one of the most likely enhancers of direct viral infection in the brain is NRP1, which is highly expressed in multiple neuron and glial cell types, including astrocytes and microglia (Cantuti-Castelvetri et al., 2020; Daly et al., 2020; Lonsdale et al., 2013)." |
| E6 (226-229) | 35858406 | SET MeSHAnatomy = "Cerebral Cortex"<br>a(TAX:"Severe acute respiratory syndrome coronavirus 2") increases r(HGNC:NRP1)<br>p(HGNC:NRP1) increases bp(GO:"viral entry into host cell") | "NRP1 is a potential candidate entry factor for infection, as it has been demonstrated to be a SARS-CoV-2 host factor (34). We observed NRP1 RNA expression in primary tissue, specifically in cortical neurons—a population, however, where we observed minimal infection (SI Appendix, Fig. 8A). When we evaluated NRP1 protein abundance, we similarly observed membrane-bound staining in neurons in the cortical plate in primary tissue, but none in dsRNA+-infected cells." |
| E7 (183-188) | 34489403 | a(TAX:"Severe acute respiratory syndrome coronavirus 2") decreases bp(GO:"maintenance of permeability of blood-brain barrier") | "To assess BBB integrity, we administered Evans blue dye (EBD) by i.p. injection in both mock-treated and SARS-CoV-2-infected hamsters at 6 dpi and evaluated its leakage into the brain, which is a classic method for evaluating BBB integrity. Compared with the mock-treated hamsters, infected animals displayed visible leakage of the EBD in the cortex (Fig. (Fig.3a).3a)." |
| E8 (297-300) | 22315722 | SET MeSHDisease = {"Alzheimer Disease", "Parkinson Disease"}<br>bp(GO:"maintenance of permeability of blood-brain barrier") neg bp(FIXME:"lymphocyte infiltration") | "Indeed, increased BBB permeability is found in both AD and PD, allowing for increased lymphocytic ingress." |
| E9 (118-120) | 32498691 | path(DO:"COVID-19") increases bp(GO:"active induction of innate immune response in host by virus")<br>path(DO:"COVID-19") increases bp(GO:"cytokine production involved in immune response") | "COVID-19 is associated with a severe innate immune response and sustained rise of systemic cytokine levels." |

| | | | |
|---|---|---|---|
| E10 (90-96) | 10381209 | a(MESH:Cytokines) -> bp(GO:"microglial cell activation") bp(GO:"microglial cell activation") -> a(CHEBI:"reactive oxygen species") SET MeSHAnatomy = "Brain" path(DO:"Alzheimer's disease") pos a(MESH:Cytokines) a(CHEBI:"anti-inflammatory drug") -\| path(DO:"Alzheimer's disease") | "Among earlier observations that stand out as pointing toward the need to consider inflammatory processes as important include: (1) Colton's demonstrations that cytokine-activated microglia produce ROS [3,4]; (2) that, in contrast to normal brain, proinflammatory cytokines are prevalent in Alzheimer's brains [5–7]; (3) the early indications that anti-inflammatory therapeutics may delay on-set of Alzheimer's disease [8]; and (4) the early demonstrations that complement and classic markers of immune-mediated damage are expressed in neurodegenerative brains [9,10]." |
| E11 (109-119) | 10381209 | bp(GO:"microglial cell activation") -> p(HGNC:IL1B) bp(GO:"microglial cell activation") -> p(HGNC:IL6) bp(GO:"microglial cell activation") -> p(HGNC:TNF) p(HGNC:IL1B) -> bp(GO:"glial cell activation") p(HGNC:IL6) -> bp(GO:"glial cell activation") p(HGNC:TNF) -> bp(GO:"glial cell activation") p(HGNC:IL1B) -> bp(GO:"astrocyte activation") p(HGNC:IL6) -> bp(GO:"astrocyte activation") p(HGNC:TNF) -> bp(GO:"astrocyte activation") bp(GO:"microglial cell activation") -> bp(GO:"inflammatory response") | "The microglia in brain are analogous to resident macrophages capable of being activated to produce cell mediators, (i.e., such as the cytokines IL-1b, IL-6, TNF-a, etc.), which in turn activate other cell populations, e.g., glia and astrocytes, to subsequently produce other cell mediators that then additionally activate microglia and other cellular populations, therefore, perpetuating and amplifying the inflammatory cascade." |
| E12 (96-100) | 22315722 | bp(GO:"microglial cell activation") increases path(HP:Neurodegeneration) bp(GO:"microglial cell activation") increases path(ADO:neuroinflammation) bp(GO:"negative regulation of adaptive immune response") increases path(HP:Neurodegeneration) bp(GO:"negative regulation of adaptive immune response") increases path(ADO:neuroinflammation) | "Aberrant proteins (e.g., modified α-synuclein) activate microglia and engage the adaptive immune system, causing neuroinflammation and neurodegeneration in PD." |
| E13 (190-192) | 30342839 | bp(GO:"inflammatory response") -> p(HGNC:SNCA, pmod(FIXME:"protein misfolding")) bp(GO:"inflammatory response") -> bp(FIXME:"alpha-synuclein aggregation") | "At the molecular level, inflammatory mediators are known to promote α-synuclein misfolding and aggregation [34]. Importantly, there are also emerging, compelling links between inflammation and the spread of α-synuclein pathology, which could explain how systemic inflammation facilitates the propagation of α-synuclein pathology from peripheral tissues to central nervous system." |
| E14 (261-267) | 22315722 | SET MeSHAnatomy = "Brain" p(HGNC:SNCA, pmod(Ub)) increases bp(GO:"Lewy body formation") p(HGNC:SNCA, pmod(Ph)) increases bp(GO:"Lewy body formation") p(HGNC:SNCA, pmod(GO:"protein oxidation")) increases bp(GO:"Lewy body formation") p(HGNC:SNCA, pmod(FIXME:"protein nitration")) increases bp(GO:"Lewy body | "These α-syn species are created by ubiquitination (Shimura et al. 2001), phosphorylation (Fujiwara et al. 2002), or oxidation and nitration (Giasson et al. 2000), and are found in LB inclusions, extraneuronally in PD brains (Lee 2008), and in the periphery of PD patients." |

| | | | |
|---|---|---|---|
| | | formation") | |
| E15 (359-368) | 22315722 | SET Species = "10090"<br>bp(FIXME:"alpha-synuclein aggregation") increases bp(GO:"microglial cell activation")<br>bp(GO:"microglial cell activation") increases a(CHEBI:"nitric oxide")<br>bp(GO:"microglial cell activation") increases a(CHEBI:superoxide) | "Aggregated α-syn activates microglia (Zhang et al. 2005), which have been shown to produce nitric oxide and superoxide in mice and inducible nitric oxide synthase (iNOS) in humans, which increases nitration of α-syn and perpetuates the proinflammatory innate immune response in PD." |
| E16 (280-295) | 22315722 | bp(GO:"microglial cell activation") increases a(CHEBI:"reactive oxygen species")<br>a(CHEBI:"reactive oxygen species") increases p(HGNC:SNCA, pmod(FIXME:"protein nitration"))<br>a(CHEBI:"reactive oxygen species") increases bp(GO:"neuron death") | "Furthermore, reactive oxygen species produced by activated microglia increases nitration of α-syn and neuronal cell death (Shavali et al. 2006); and in turn, immune T cells that recognize nitrated α-syn (N-α-syn) enhance the neurotoxic activities of microglia in the acute 1-methyl-4-phenyl-1,2,3,6-tetrahydropyridine (MPTP) mouse model of nigrostriatal degeneration . Activated T cells and B cells are then able to enter the CNS more readily and migrate to the site of neuronal injury." |
| E17 (97-106) | 24275605 | SET Species = "9606"<br>SET MeSHAnatomy = "Neurons"<br>path(DO:"Parkinson's disease") increases path(ADO:neuroinflammation)<br>SET MeSHAnatomy = "Substantia Nigra"<br>SET Anatomy = "midbrain"<br>path(DO:"Parkinson's disease") increases a(MESH:"Lewy Bodies")<br>path(DO:"Parkinson's disease") decreases a(MESH:"Dopaminergic Neurons") | "PD pathology is characterized by chronic neuroinflammation, Lewy body inclusions, and loss of dopamine-producing (DA) neurons in the substantia nigra parts compacta (SNpc) of the midbrain [4, 6]" |
| E18 (233-238) | 37458167 | p(HGNC:APOE, var("?")) -> path(DO:"Alzheimer's disease")<br>SET Disease_Severity = "severe"<br>p(HGNC:APOE, var("?")) -> path(DO:"COVID-19")<br>bp(GO:aging) -- path(DO:"COVID-19") | “Indeed, carrying the Alzheimer’s disease risk allele APOE ε4 (Kuo et al., 2020; Wang et al., 2021c) or being of an advanced age (González-García et al., 2021) increase the risk of severe COVID-19.” |
